# Functional specialization of mPFC-BLA and mPFC-NAc pathways in affective state representation

**DOI:** 10.1101/2024.05.28.596238

**Authors:** Chien-Hsien Lai, Gyeongah Park, Pan Xu, Xiaoqian Sun, Qian Ge, Zhen Jin, Sarah Betts, Xiaojie Liu, Qingsong Liu, Rahul Simha, Chen Zeng, Hui Lu, Jianyang Du

## Abstract

Effective emotional processing, crucial for adaptive behavior, is mediated by the medial prefrontal cortex (mPFC) via connections to the basolateral amygdala (BLA) and nucleus accumbens (NAc), traditionally considered functionally similar in modulating reward and aversion responses. However, the functional specialization of the mPFC→BLA and mPFC→NAc pathways in representing affective states remains unclear. We found that while overall firing patterns appeared consistent across emotional states, deeper analysis revealed distinct variabilities. Specifically, mPFC→BLA neurons, especially “center-ON” neurons, exhibited heightened activity during behaviors classically associated with anxiety-like states, suggesting their involvement in aversive behavioral regulation. Conversely, mPFC→NAc neurons were more active during exploratory and approach-related behaviors, implicating them in the processing of positively valenced behavioral states. Notably, mPFC→NAc neurons showed significant pattern decorrelation during social interactions, suggesting a pivotal role in processing social preference. Additionally, repeated win/loss outcomes in the tube test produced distinct hierarchy-dependent behavioral changes and elevated corticosterone levels in loser mice, supporting the biological relevance of these behaviorally defined states. Together, these findings reveal pathway-specific representations of affect-related behavioral states in mPFC circuits and provide a framework for understanding how prefrontal outputs organize adaptive behavior across environmental contexts.

## INTRODUCTION

Recent advancements in neuroscience have significantly advanced our understanding of the complex roles played by the medial prefrontal cortex (mPFC) pathways to the basolateral amygdala (BLA) and nucleus accumbens (NAc). These pathways are pivotal in emotional and motivational processing, mediating behaviors that involve assessing conflicts between reward and aversion, thus enabling the brain to adaptively respond to diverse environmental stimuli (*1, 2*).

Both the mPFC→BLA and mPFC→NAc pathways play key roles in shaping emotional responses and guiding decisions when animals face potential rewards or threats (*3*). These circuits adjust behavior when reward seeking must be suppressed in the presence of aversive cues, thereby maintaining a balance between risk and reward in emotionally charged contexts. The prelimbic (PL) subdivision of the mPFC sends excitatory projections to the BLA, a pathway causally linked to fear and anxiety: optogenetic activation of PL→BLA maintains fear memories and induces anxiety-like behaviors (*4, 5*). In contrast, the mPFC→NAc pathway is implicated in exploration, reward-seeking, and motivational drive, with optogenetic activation of PL→NAc projections biasing exploratory behavior and social approach (*6*).

The mPFC-BLA circuit also integrates reward-aversion conflicts. Ishikawa et al. showed that infralimbic (IL) projections to the BLA are required to suppress reward-seeking when paired with shock punishment (*7*), an effect likely mediated through BLA influence on the NAc shell (*8*). Inactivation of PL, BLA, or NAc shell disrupts avoidance (*9, 10*). PL projections to BLA and NAc have opposing effects on avoidance: PL-BLA activation facilitates avoidance, while PL-NAc activation inhibits it (*11*). During avoidance retrieval, PL neurons projecting to BLA are activated, while avoidance extinction activates both PL-NAc and IL-NAc projections (*12*). Finally, Kim et al. reported that activation of mPFC→NAc lateral shell neurons engaged by aversive stimuli can suppress reward seeking, underscoring the role of this pathway in weighing competing motivational demands (*13*).

Together, these studies highlight both shared and distinct roles of the mPFC→BLA and mPFC→NAc pathways in shaping emotional and motivational behaviors. While both circuits modulate responses to reward-threat conflict, they do so through partially divergent mechanisms, emphasizing the complexity of prefrontal control over affect and action. Despite substantial progress, however, many questions about the precise neural mechanisms within these pathways remain unresolved. Traditional approaches, including pharmacological manipulations, c-fos mapping, circuit tracing, and even pathway-specific optogenetics, have provided critical insights but lack the resolution to selectively manipulate emotion-related ensembles. Because these techniques often do not distinguish between specific cell types or functionally defined subpopulations, important circuit- and cell-level differences may remain obscured, limiting our understanding of how these pathways implement their distinct functions.

Building on this foundation, we employed in vivo imaging to examine the network dynamics of the mPFC→BLA and mPFC→NAc pathways during emotionally salient behaviors. Our goal was to directly compare how these circuits encode affective states, focusing on the similarities and differences in their neuronal activation patterns and ensemble dynamics. Because a key challenge in such analyses is determining whether behaviorally defined states reflect meaningful internal conditions, we additionally employed a repeated social competition paradigm to assess whether these states are associated with measurable physiological and behavioral changes. This complementary approach allowed us to identify pathway-specific activity patterns while independently validating the biological relevance of the behavioral states used in our analyses.

## RESULTS

### Differential activity patterns of mPFC→NAc and mPFC→BLA neurons during anxiety-like and exploratory behavioral states

To distinguish the mPFC→BLA and mPFC→NAc neurons, we employed retrograde AAV viruses carrying genes encoding the Ca^2+^ indicator GCaMP6m, delivering them to the BLA and NAc of adult female wildtype mice at the age of ∼20 weeks, respectively (**Fig. 1A**). This allowed for the infection of mPFC neurons projecting to either of these regions, leading to GCaMP6 expression in the soma situated in the mPFC (**Fig. 1B**). In vivo Ca^2+^ imaging was conducted on mPFC→BLA and mPFC→NAc neurons using a portable miniaturized microscope (**Fig. 1C**). The activity of individual neurons was continuously monitored throughout the behavioral assays. We synchronized the Ca^2+^ imaging with the behavioral recordings, following the same neurons within each mouse across the different test sessions (see Methods).

**Figure 1.**
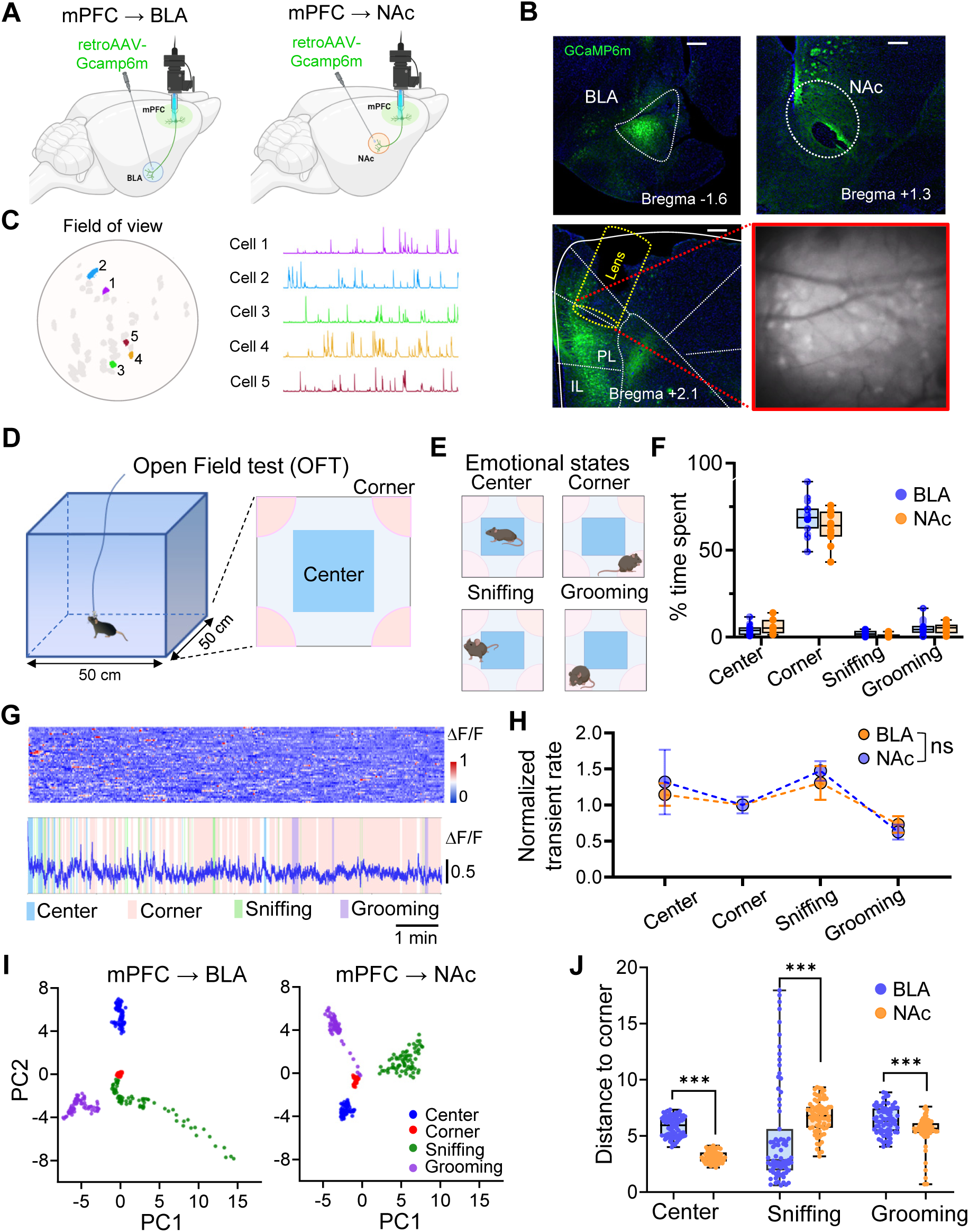
The mPFC→NAc and mPFC→BLA neurons demonstrated distinct activity patterns across various emotional states. (A) Schematic showing retrograde AAV-GCaMP6 injection in the NAc and BLA, respectively. In vivo calcium imaging was performed in the mPFC neurons expressing GCaMP6 via miniscope. (B) Top: An example of AAVretro-GCaMP6 injection and expression in the BLA and NAc, respectively. Bottom: An example of GCaMP6m expression in the mPFC with AAVretro-GCaMP6 injection in the NAc and an image of Ca2+ fluorescence recorded with miniaturized microscope. The 1-mm GRIN lens covered the full depth of the prelimbic cortex. Green, GCaMP6m. Scale bars, 200 mm. (C) Left: the field of view under a GRIN lens in one mouse with identified neurons numbered and colored. Right: fluorescence traces of example neurons marked in the above panel. (D). Schematic of the open field test (OFT) in an open 50 cm X 50 cm arena. The pink areas represent corners and the blue area represents the center. (E) Illustrations depicting typical mouse behaviors associated with different emotional states observed during the OFT: Center (exploration and higher anxiety), Corner (lower anxiety), sniffing (sensory exploration and vigilance, possibly indicating curiosity, cautious exploration, or heightened alertness), and grooming (stress relief or comfort). (F) The percentage of time that mice spent in the center, corners, sniffing and grooming during 10-min open field test. n = 15 mice for mPFC→BLA group; n = 11 mice for mPFC→NAc group. The time-allocation pattern across behaviors is similar for Amy and NAc cohorts. A two-way mixed-effects model (Region × Behavior) showed a strong main effect of Behavior (Corner » Center; Sniffing < Center), but no main effect of Region and no Region × Behavior interaction (all P > 0.1). (G) Heatmap and time-series plot of neuronal activity (ΔF/F) across the test duration. The heatmap above shows the fluctuation in activity level across neurons, while the plot below aligns the averaged fluctuations of these neurons with observed behaviors (Center, Corner, Sniffing, Grooming) within the arena. (H) Normalized transient rates of neuronal activity during different behaviors. The mixed-design ANOVA revealed no main effect of region (P = 0.14) and no significant Region × Behavior interaction (P > 0.3), indicating that mPFC→BLA and mPFC→NAc neurons exhibited similar behavioral modulation patterns. (I) PCA plots for mPFC→BLA and mPFC→NAc pathways illustrating the distribution of neuronal activity patterns for different observed behaviors (Center: blue, Corner: red, Sniffing: green, Grooming: purple). Each point represents an individual neuronal recording. Quantitative analysis confirmed that neural population activity during corner behavior was highly similar between mPFC→BLA and mPFC→NAc pathways (Pearson r = 0.64, p < 1 × 10⁻⁹), consistent with comparable encoding of this low-anxiety state. (J) Summary of the distances of neuronal activity clusters from the center of Corner behavior in the OFT for both pathways. distances were computed using PC1 and PC2 coordinates. ***P < 0.001, Mann-Whitney U test.

We first performed the open field test (OFT), a widely employed behavioral assay in rodent research for assessing exploratory and anxiety-like behaviors (*14*) (**Fig. 1D**). This test leverages the inherent conflict rodents experience between their desire to explore new environments and their instinctive aversion to exposed spaces. We monitored and recorded the mouse’s movements and actions over a 10-minute period, specifically tracking time spent in the center versus the corner, wall-sniffing, and grooming. These behaviors observed during the open field test, such as spending time in corners, the center area, grooming, and sniffing walls (**Fig. 1E**), offer valuable insights into the mice’s affectively salient behavioral states. Typically, corner occupancy is commonly interpreted as reflecting a relatively safer or less exposed state. Conversely, exposure to the center area may induce higher anxiety due to increased exposure and reduced safety compared to the arena’s edges or corners. Grooming behaviors can signal stress regulation or self-directed coping processes, depending on context, while wall-sniffing reflects sensory exploration and vigilance, possibly indicating curiosity, cautious exploration, or heightened alertness (*14, 15*). Our findings demonstrate that mice consistently spend significant time in corners during the 10-minute session. Additionally, they allocate around 10% of the time to the center area or grooming, and approximately 5% of the time to wall-sniffing (**Fig. 1F**). To ensure accurate behavioral categorization, “sniffing” was defined as episodes in which the mouse’s nose was within ∼1 cm of the wall and moving in small scanning motions without locomotion for ≥1 s, as detected by the TopScan system. Sniffing events occurring within the corner zones were excluded from the corner category to maintain mutual exclusivity between the two behaviors. Accordingly, the corner time in **Fig. 1F** does not include sniffing episodes, allowing for a clear distinction between directed exploratory (sniffing) and stationary, anxiety-associated (corner) states.

To determine whether the pattern of neuronal activities in the mPFC→BLA and mPFC→NAc pathways differs in response to distinct affective behavioral states, we first estimated the Ca^2+^ transient rate. During our observation of mPFC→BLA and mPFC→NAc neuron activity (**Fig. 1G**) during the open field test, we determined that the collective averaged Calcium transient rate of the neurons in these two pathways did not exhibit discernible differences while the mice were situated in the center, corner, or during sniffing or grooming (**Fig. 1H** and **Supplementary Fig. 1**). To delve deeper, we conducted a Principal Component Analysis (PCA) to assess the population activity pattern linked to various behaviors, encompassing center or corner dwelling, grooming, and sniffing. The PCA plots in **Fig. 1I** reveal how neuronal activities associated with various behaviors differ between these pathways, with each point representing recorded neuronal activity during specific behaviors projected onto the first two principal components (PC1 and PC2). These components capture the most significant variances within the dataset. These two plots display distinct clustering of activity patterns with clear separations among the different behaviors, particularly between center and corner behaviors, as well as sniffing and grooming. To account for the apparent spread in the mPFC→BLA trajectories during sniffing, we examined the temporal progression of neural activity and found that early frames clustered tightly, whereas later frames diverged along PC1, reflecting transitions into other behaviors. Shortening the analysis window to 2-3 s post-onset reduced this spread, confirming that the variance observed in the 5 s window arises from within-bout behavioral transitions rather than overlap with corner activity or time-in-arena effects.

To quantify the similarity of behavioral representations between the two projection pathways, we performed a canonical correlation analysis (CCA) on the reduced PCA representations of averaged behavioral bouts. Corner behavior exhibited the smallest cross-pathway difference (mean absolute difference = 0.12, p = 0.00039 < 0.001), indicating highly overlapping representations between mPFC→BLA and mPFC→NAc neurons. In contrast, Center (0.47, p = 0.968), Sniffing (0.43, p = 0.0002), and Grooming (0.34, p = 0.0002) showed substantially larger differences. These findings confirm that corner activity is encoded most similarly across the two pathways, supporting its use as a stable, low-anxiety reference state for comparison.

We next quantified distances from Corner behavior to other behavioral states across both pathways (**Fig. 1J**). Distances between behavioral clusters were calculated in the PC1-PC2 space as Euclidean distances from the centroid of the corner distribution, which served as a reference state because mice spent the majority of time in the corners. While both pathways exhibited similar neural patterns for Corner behavior, Center and Grooming states were markedly farther from Corner in the mPFC→BLA pathway, indicating more distinct neural encoding of anxiety-related versus non-anxiety behaviors. This differentiation was especially pronounced for Grooming, suggesting that the mPFC→BLA pathway plays a key role in distinguishing stress- and coping-related states. Conversely, the mPFC→NAc pathway showed stronger separation between Sniffing and Corner behaviors, highlighting its selective involvement in differentiating exploratory behaviors from those associated with anxiety (**Fig. 1I, J**).

To ensure that the observed distance relationships were not driven by random label structure or temporal correlations, we implemented permutation-based null controls. Behavioral labels were shuffled within each mouse and session, and circular time-shifts were applied to each neuron within bouts to preserve autocorrelation. The observed Corner distances and between-pathway differences exceeded the 95th percentile of both null distributions and remained significant after 10,000 permutations (p_perm < 0.01). These controls confirm that the reported cluster relationships reflect genuine structure in neural population activity. Together, these analyses underscore the distinct functional roles of the two mPFC projection pathways: the mPFC→BLA pathway preferentially encodes anxiety-related versus non-anxiety states, whereas the mPFC→ NAc pathway more selectively distinguishes exploratory from anxiety-like behaviors.

The above differentiation in population activity was not revealed by averaging the transient rate. When considering averaging the transient rate across a population can potentially mask characteristics unique to neurons that represent the emotional states of mice associated with these behaviors. We thus narrowed our focus to neurons that exhibited increased activity during heightened anxiety moments, such as entering the center zone. Termed “center-ON” neurons, these neurons displayed substantially increase in activity surrounding the mouse’s entry into the center zone, suggesting sensitivity to a behavioral shift from a low- to high-anxiety-like state triggered by entry into the anxiogenic center area (**Fig. 2A**). This increase in activity of the center-ON neurons in the mPFC→BLA pathway was not observed when the mice entered corner zones or engaged in sniffing or grooming (**Fig. 2B**). Upon comparing the percentage of center-ON neurons in the two pathways, we observed a higher prevalence of center-ON neurons in the mPFC→BLA group compared to the mPFC→NAc group (**Fig. 2C** and **Supplementary Fig. 2**). This finding supports the earlier observation that neurons in the mPFC→BLA pathway exhibit more pronounced encoding of anxiety-like states compared to those in the mPFC→NAc pathway. The distribution of center-ON neurons within the mPFC→BLA and mPFC→NAc pathways was consistent across animals and spanned both superficial and deep layers, as the 1-mm GRIN lens encompassed the full depth of the prelimbic cortex, indicating that their activity was not layer-specific. When we projected the activity patterns of these center-ON neurons onto the spatial distribution of the open field arena (**Fig. 2D**), both sets of center-ON neurons displayed heightened activity in the center zone. However, the center-ON neurons in the mPFC→NAc population exhibited increased activity during sniffing as well, which was not observed in the mPFC→BLA population (**Fig. 2E** and **Supplementary Fig. 3**). This finding implies that these two sets of center-ON neurons serve distinct roles in encoding behaviorally defined affective states in mice. Specifically, the mPFC→NAc neurons appear to be more involved in positively valenced behavioral states associated with exploration, while the mPFC→BLA neurons are more active during negatively valenced states linked to anxiety-like behavior. To reinforce this notion, we conducted the PCA exclusively on the center-ON neuron populations (**Fig. 2F**). Intriguingly, PCA scatter plots reveal behavioral clustering and spatial separations, demonstrating that different behaviors influence neuronal engagement differently in these two pathways. Again, we quantified the distances of neuronal activity patterns associated with Center, Sniffing, and Grooming from Corner behavior in both pathways (**Fig. 2G**). The result showed significant differences in the encoding of behaviorally defined affective states by center-ON ensembles in the two pathways. Similar to **Fig. 1J**, the greater separation between Center and Corner behaviors in the mPFC→BLA pathway suggests that this projection may play a critical role in modulating stress- and anxiety-related responses, underscoring its potential relevance as a therapeutic target for anxiety disorders. In contrast, the mPFC→NAc pathway appears more broadly engaged in general exploratory behaviors and less specialized in processing anxiety-specific signals (**Fig. 2F and G**). The presence of distinct center-ON neuronal populations further highlights the functional divergence between these pathways, with the mPFC→BLA and mPFC→NAc circuits exhibiting pathway-specific encoding of aversive versus exploratory behavioral states.

**Figure 2.**
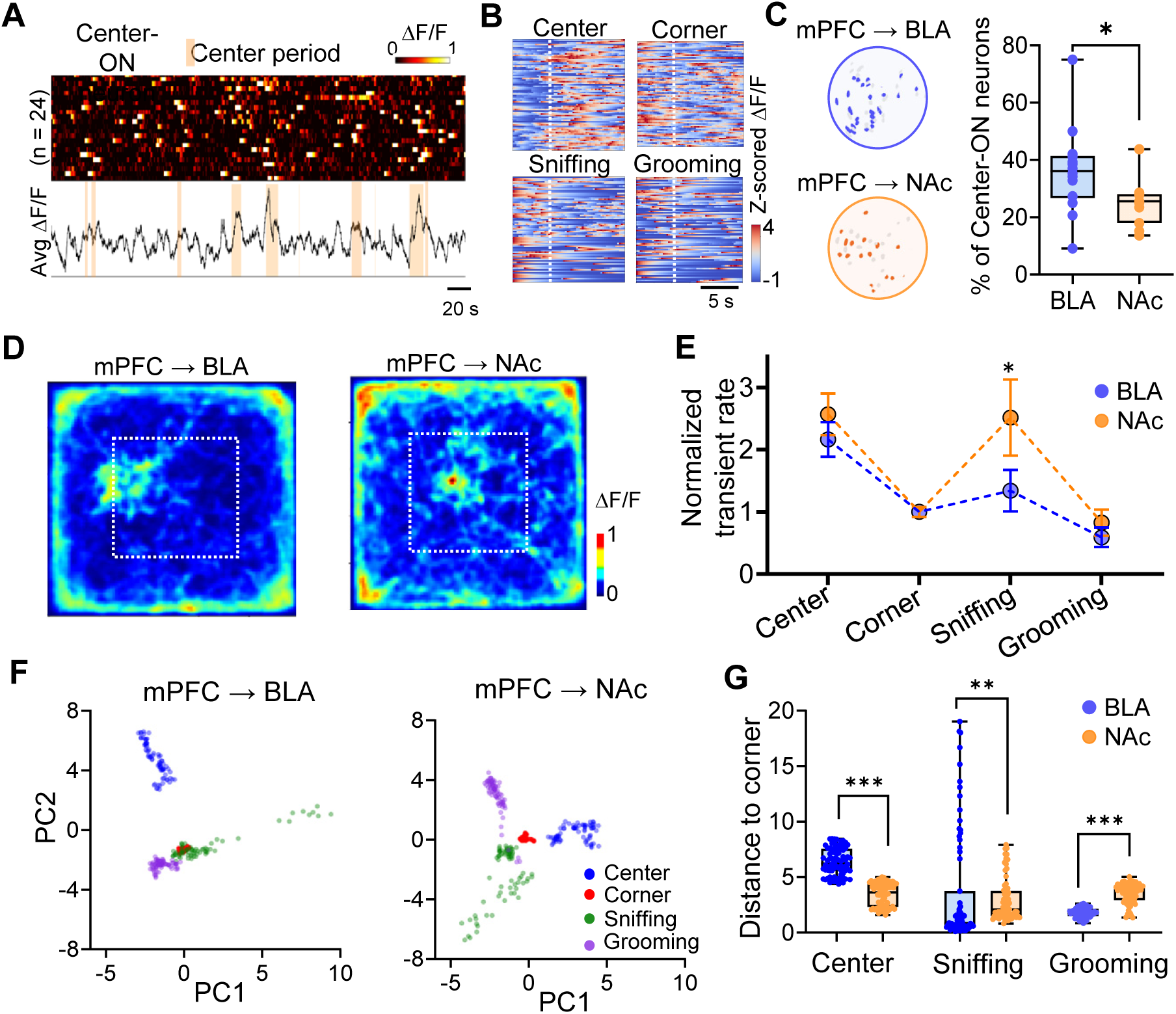
Distinct encoding of emotional status by center-ON neurons in the mPFC pathways. (A) Ca^2+^ traces of the averaged activity of center-ON neuron ensembles around the onset of center entry (5 s before to 5 s after). The total number of 24 center-ON neurons (n = 24) recorded in a mouse of the mPFC→BLA groups that used to calculate the representative averaged trace. Solid lines represent the averaged value. (B) Representative heatmaps depicting the activity patterns of mPFC→BLA neurons across four different behavioral contexts: Center, Corner, Sniffing, and Grooming. Each column corresponds to a different behavior, with the intensity of color indicating the level of neuronal activity. (C) Left: spatial distributions of center-ON neurons among the mPFC→BLA and mPFC→NAc neurons in one representative mouse from each group. Right: quantification of the percentage of center-ON neurons showed that the BLA pathway contained a significantly higher proportion of center-ON neurons compared to the NAc pathway. * P < 0.05, Mann-Whitney U test (Mann–Whitney U = 47, P = 0.049). (D) Spatial heatmaps of neuronal activity across the open field arena of all observed center-ON neurons in one example mouse of mPFC→BLA and mPFC→NAc groups. The color bar indicates the averaged normalized z-score. (E) Normalized transient rate of neuronal activity across behavioral states for mPFC→BLA and mPFC→NAc neurons. A two-way mixed-design ANOVA revealed a significant Region × Behavior interaction (P = 0.048), indicating that mPFC→NAc neurons exhibited higher activity during sniffing compared to mPFC→BLA neurons, while no regional differences were found in center, corner, or grooming states. * P < 0.05; two-way mixed-design ANOVA followed by Tukey’s post hoc test. (F) PCA plots illustrating the distribution of neuronal activity patterns during different behaviors for mPFC→BLA and mPFC→NAc pathways. Points are color-coded by behavior type (Center: blue, Corner: red, Sniffing: green, Grooming: purple). (G) Summary of the distances from the center of Corner behavior in terms of neuronal activity for each behavior in both mPFC pathways. ** P < 0.01, *** P < 0.001, Mann-Whitney U test.

### Distinct activation of center-ON Neurons during anxiety-like behavioral states

To further validate our previous findings, we conducted the Elevated Plus Maze (EPM) test with the same mice one day following the OFT, as shown in **Fig. 3A**. The EPM test assesses anxiety by noting whether mice explore the exposed and elevated open arms (indicative of higher anxiety) or prefer the sheltered, enclosed arms (indicative of lower anxiety) (*16*). Our observations revealed that mice predominantly stayed in the closed arms, particularly favoring one closed arm (preferred closed arm, closed-P) over the other (non-preferred closed arm, closed-NP), suggesting a strong preference for a safer environment, despite both closed arms being designed to offer security (**Fig. 3B**). Neuronal activity in the mPFC→BLA and mPFC→NAc pathways was recorded during the EPM test. The neurons exhibited increased transient rates in the open arms, indicative of an anxiety-like behavioral response, though no significant differences were observed in the overall activity patterns between these pathways (**Fig. 3C and Supplementary Fig. 4**). This is consistent with earlier results from the OFT (**Fig. 1H**). Subsequent analyses focused on “center-ON” neurons previously identified during the OFT, examining their responses across different behavioral states in the EPM (**Fig. 3D and E**). Notably, center-ON neurons in the mPFC→BLA pathway demonstrated significantly heightened activity in the open arms compared to the closed arms, particularly when mice entered the open arms (**Fig. 3F**). The calcium transient rates for these neurons were markedly higher than those in the mPFC→NAc pathway, especially when mice were in the less familiar open-NP arm, suggesting that the unfamiliarity of this arm enhances anxiety-like behavior, and the mPFC→BLA pathway is particularly sensitive to this affective shift (**Fig. 3G and Supplementary Fig. 5**). These findings corroborate our initial observations from the OFT, highlighting the critical role of mPFC→BLA neurons in modulating responses to anxiety-like behavioral states in mice. The pronounced activity differences in these neurons suggest their heightened sensitivity to anxiogenic stimuli and their potential involvement in perceiving and processing such states. Together, the results indicate that center-ON neurons in the mPFC→BLA pathway may play a more prominent role in detecting and potentially regulating anxiety-like behaviors compared to those in the mPFC→NAc pathway.

**Figure 3.**
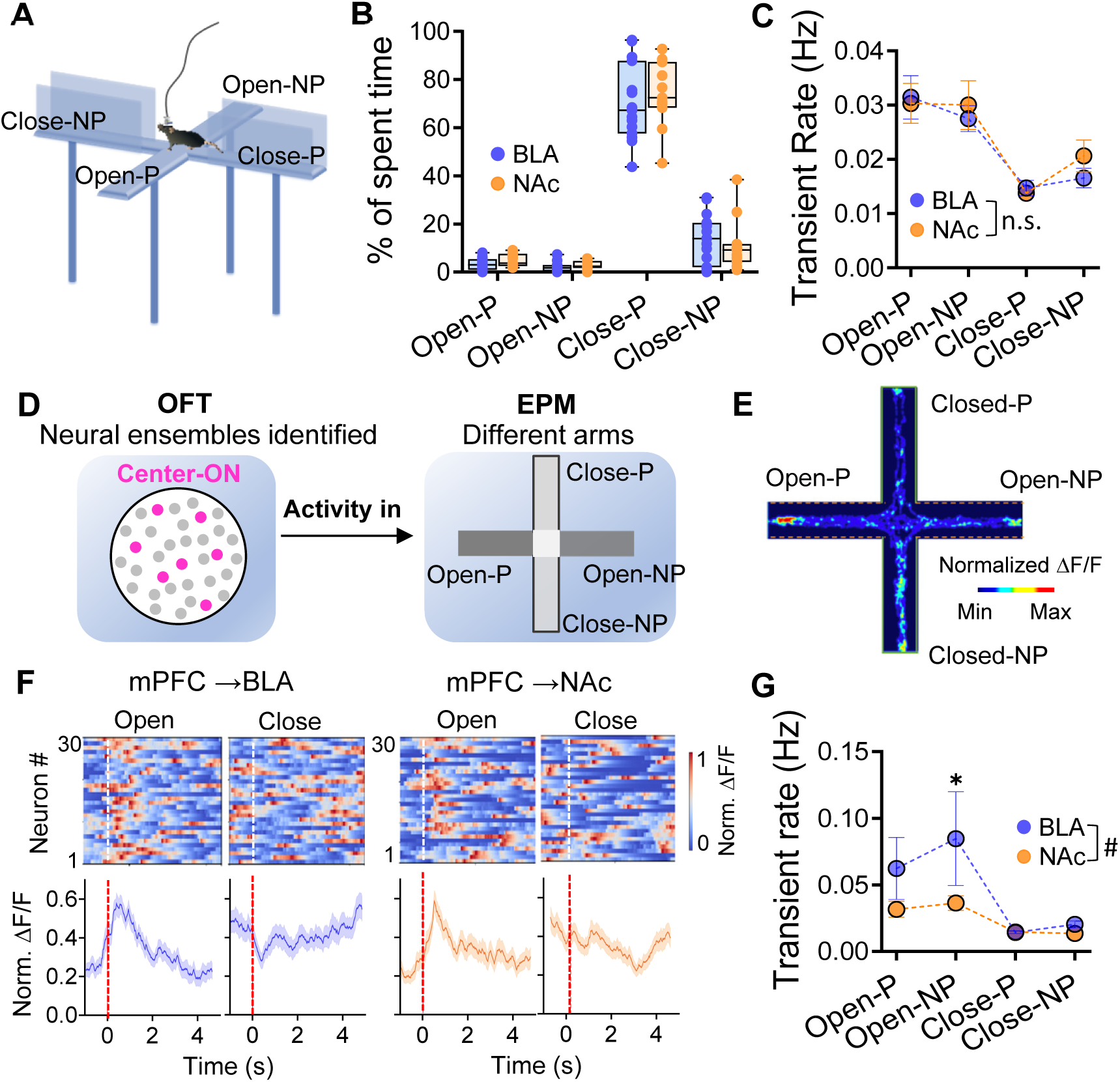
The mPFC→NAc and mPFC→BLA neurons demonstrated distinct activity patterns across various emotional states during EPM test. (A) Schematic illustration of the Elevated Plus Maze (EPM) test setup showing the layout of open (Open-P, Open-NP) and closed (Close-P, Close-NP) arms used to assess anxiety-related behaviors in mice. (B) Box plots displaying the percentage of time spent by mice in the different arms of the EPM (Open-P, Open-NP, Close-P, Close-NP) for both mPFC→BLA and mPFC→NAc pathways. Data indicate variations in time spent across different arms, highlighting behavioral preferences. (C) Averaged transient rates (Hz) of neuronal activity of all recorded neurons in the mPFC→BLA and mPFC→NAc pathways in each arm of the EPM. Data are represented as mean ± SEM. Mixed-effects analysis (Region × Arm) showed higher transient rates in open arms relative to closed arms (Open-P vs Close-NP: β=0.0149, P=7.3×10⁻⁶; Open-NP vs Close-NP: β=0.0107, P=0.0019) with no overall Region effect (p=0.304) and no Region × Arm interaction (all P>0.29), indicating the activity pattern of the mPFC→BLA and mPFC→NAc pathways in each arm of the EPM is similar. (D) Analysis design of observing the neural activities of center-ON ensembles of OFT in different EPM arms. (E) Mean Ca2+ signal across the EPM arms of all observed center-ON neurons in one example mouse. The color bar indicates the averaged normalized z-score. (F) Example 30 center-ON neurons (top) and corresponding averaged activity (bottom) around the onset of arm entry during the EPM test. Time 0 represents the onset of arm entry. (G) Group data showing the mean transient rate of center-ON neurons across four EPM arms for mPFC→BLA and mPFC→NAc pathways. A two-way mixed-effects ANOVA with Region (BLA vs NAc) as a between-subject factor and Condition (Opent-P, Open-NP, Close-P, Close-NP) as a within-subject factor revealed a significant Region × Condition interaction (F(3,72) = 3.21, P = 0.030).Tukey’s post hoc tests showed that BLA exhibited significantly higher transient rates than NAc during the Open-NP condition (P = 0.018), while no significant regional differences were found in other three arms (P > 0.1). Data are represented as mean ± SEM. n = 9–13 mice for mPFC→BLA; n = 8-11 mice for mPFC→NAc.

### The mPFC→NAc neurons, as opposed to mPFC→BLA neurons, encode emotional information associated with social preference

Social behavior can serve as an effective indicator of diverse affective states in individuals (*17*) and is encoded by the mPFC neuron ensembles with distinct activity patterns (*18*). We thus investigated whether distinct activity patterns are exhibited by mPFC→BLA and mPFC→NAc neurons, correlating with different socially relevant behavioral states. The social test was conducted using a three-chamber apparatus (**Fig. 4A**). The two end chambers were equipped with either another mouse or an object. The subject mouse underwent a 10-minute acclimation phase in the central 45 cm chamber, with the 10 cm end compartments empty. Subsequently, three ten-minute testing sessions (S1, S2, and S3) were conducted, each involving different stimuli in the end chambers to assess interaction. Utilizing automated tracking software, the time spent within the “sniffing zone” was measured. Notably, the mice exhibited a preference for interacting with fellow mice over objects in S1 and S2. In S2, this preference gap narrowed due to the relocation of objects. During S3, when the choice was between two mice, the test subjects demonstrated a significant preference for exploring the novel mouse (**Fig. 4B**), consistent with established patterns of social novelty behavior. These findings collectively signify a normal level of sociability as observed in the three-chamber test.

**Figure 4.**
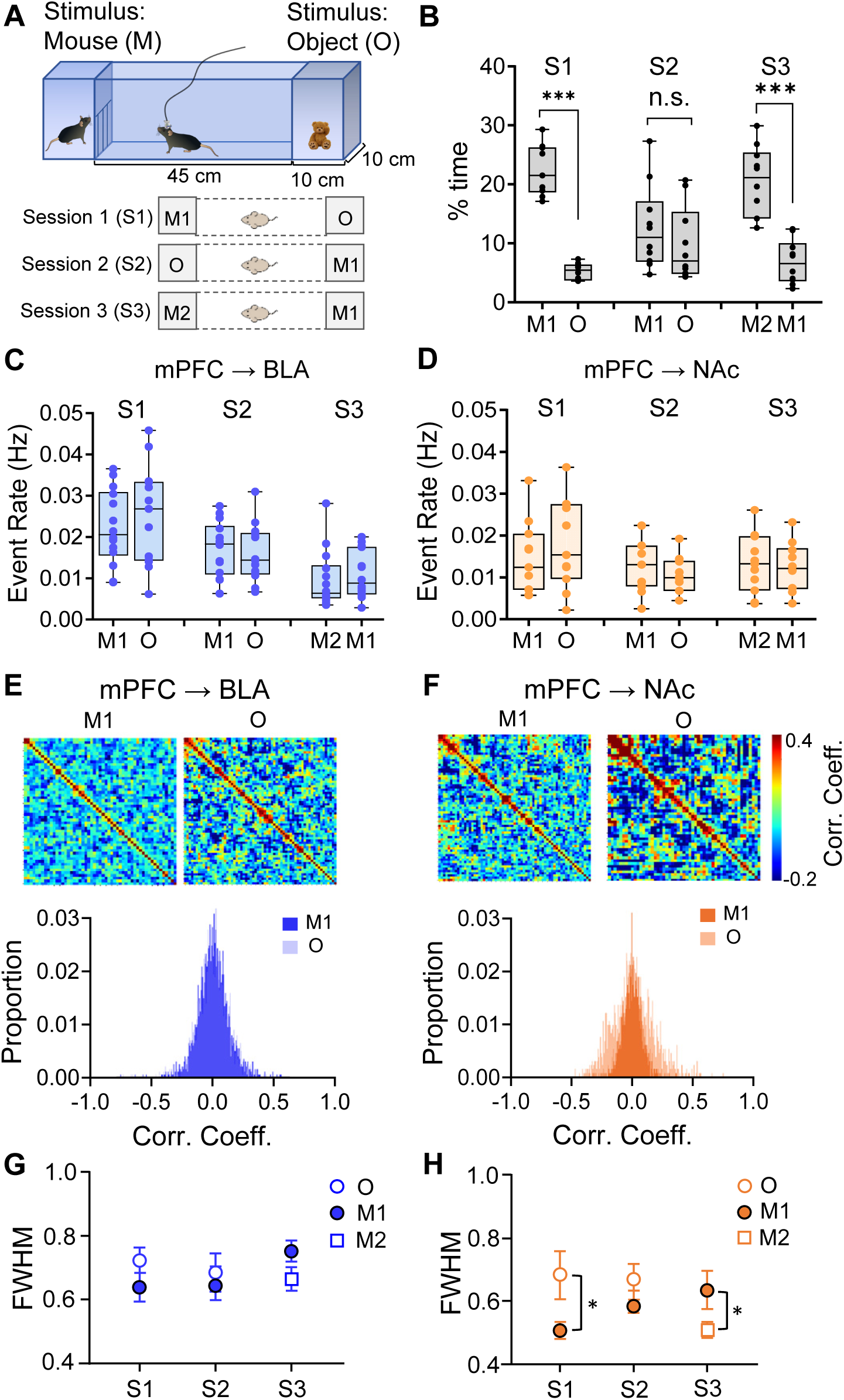
Pattern decorrelation is shown by the mPFC→NAc neurons, but not mPFC→BLA neurons. (A). Top: the apparatus for the social interaction test. The subject mouse is in the middle 45 cm-long chamber; the 10 cm end compartments contain different stimuli. Bottom: the three sessions of the social interaction test. In the first 10-minute test session (S1), a strange mouse (M1) and an object (O) were placed in the end chambers; session 2 (S2) used the same stimuli but swapped their positions; in session 3 (S3) a new mouse (M2) replaced O, so that the subject mouse must choose whether to interact with a familiar or strange mouse (M1 vs. M2). (B). The percentage of time that mice (n = 10) spent interacting with stimuli in each session. Data are shown as mean ± SEM. Paired t-test, S1: t(9) = 12.71, P = 1.2×10⁻⁶, Cohen’s d = 4.02; S2: t(9) = 0.88, P = 0.40, Cohen’s d = 0.28; S3: t(9) = 7.29, P = 6.7×10⁻^5^, Cohen’s d = 2.31. Mice spent significantly more time with the social mouse (M1) than with the object (O) in S1 or the new social mouse (M2) than with the old social mouse (M1) in S1. (C). The averaged calcium event rate of mPFC→BLA group when mice were engaged with different stimuli over the three sessions. n = 13 mice. Data are represented as mean ± SEM. Paired two tailed t-test, S1: t(12) = -0.77, P = 0.46, Cohen’s d = -0.22; S2: t(12) = 033., P = 0.74, Cohen’s d = 0.09; S3: t(12) = -0.39, P = 0.070, Cohen’s d = - 0.11. The transient rate of the mPFC→BLA neurons showed no significant difference during the interactions with the two stimuli in each session. (D). The averaged calcium event rate of mPFC→NAc group when mice were engaged with different stimuli over the three sessions. n = 11 mice. Data are represented as mean ± SEM. Paired two tailed t-test, S1: t(10) = -0.88, P = 0.40, Cohen’s d = -0.27; S2: t(10) = 178, P = 0.11, Cohen’s d = 0.56; S3: t(10) = -1.45, P = 0.18, Cohen’s d = 0.46. The transient rate of the mPFC→NAc neurons showed no significant difference during the interactions with the two stimuli in each session. (E-F). Top: raster plots of correlation coefficient of paired mPFC neurons during interactions with different stimuli, from a representative mouse in mPFC→BLA and mPFC→NAc group, respectively. Bottom: distribution of pair-wise Pearson correlation coefficients among these recorded mPFC neurons in responding to stimuli in each session. The blue and orange plots represent M1 interactions, while the lighter colors represent the interaction with the other stimulus (O). (G). The averaged full width at half maximum (FWHM) of correlation coefficient distribution of individual mice in mPFC→BLA (n = 8) group during interactions with different stimuli Data are shown as mean ± SEM. Paired two tailed t-test, S1: t(7) = -1.63, P = 0.15, Cohen’s d = -0.58; S2: t(7) = -0.41, P = 0.69, Cohen’s d = -0.14; S3: t(7) = -1.63, P = 0. 15, Cohen’s d = 0.57. The correlation coefficient of the mPFC→BLA neurons showed no significant difference between two stimuli. (H). The averaged full width at half maximum (FWHM) of correlation coefficient distribution of individual mice in mPFC→NAc (n = 8) group during interactions with different stimuli Data are shown as mean ± SEM. *P<0.05, Paired two tailed t-test, S1: t(7) = -2.75, P = 0.025, Cohen’s d = -0.92; S2: t(7) = -0.92, P = 0.39, Cohen’s d = - 0.33; S3: t(7) = -2.87, P = 0.021, Cohen’s d = -0.96. The correlation coefficient of the mPFC→NAc neurons showed significantly stronger separation between the two stimuli during S1 and S3 than during S2.

To explore the neuronal activity associated with distinct social states, we closely monitored the excitatory mPFC→BLA and mPFC→NAc neurons within the prelimbic region of the mPFC. Surprisingly, both the mPFC→BLA and mPFC→NAc neurons exhibited little discernible difference in their Ca^2+^ transient rate while the mice interacted with the social stimulus (mouse 1, M1) and the nonsocial stimulus (object, O) (**Fig. 4C and D**). Likewise, no significant variation was observed during S3. Given that interactions with social versus nonsocial stimuli are known to elicit distinct affective states, as reflected in social preference behavior, our findings suggest that the neural encoding of these states is not solely dependent on the overall firing rate of mPFC→BLA and mPFC→NAc neurons.

Next, we analyzed the ability of these two groups of neurons to convey information through coactivity patterns (*19*). Functional correlations (coactivity) within the circuit can be identified using Pearson correlation coefficients, and pattern decorrelation serves to clarify overlapping activity patterns. This mechanism is akin to how the olfactory bulb distinguishes closely related odorants based on minor structural variations (*20*). In a previous study, we demonstrated that pattern decorrelation of mPFC excitatory circuit neuronal activities is indispensable for social preference (*21*). This led us to contemplate whether differences existed between the mPFC→BLA and mPFC→NAc neurons. We thus calculated the pairwise Pearson correlation coefficients of neuronal activities specifically during the interaction with the social or object stimulus (**Fig. 4E and F**). The pairwise correlation coefficients for different stimuli across the recorded mPFC→BLA and mPFC→NAc neurons from all mice exhibited a distribution around zero, albeit with varying widths. Intriguingly, we observed a significant pattern decorrelation only within the mPFC→NAc neurons, evident by the full-width at half maximum (FWHM) of the correlation distribution being lower for the more-attractive stimuli in each session (**Fig. 4F and H**). In contrast, such decorrelation was not observed in the mPFC→BLA neurons (**Fig. 4E and G**). The broader tuning profile of mPFC→NAc neurons suggests a more sustained and robust encoding of social preference, reflected in distinct coactivity patterns across ensembles. While this enhanced encoding aligns with the positive valence often associated with social interactions, it does not by itself establish direct encoding of emotional states. Instead, these findings indicate that the mPFC→NAc pathway may play a prominent role in processing and guiding social behavior, whereas mPFC→BLA neurons appear less engaged in differentiating between social and nonsocial stimuli at the level of ensemble coactivity.

We next explored the distinct activity patterns of mPFC→BLA and mPFC→NAc neurons during social (M1) and nonsocial (object, O) interactions by comparing all recorded neurons and the subset of center-ON neurons previously identified (**Fig. 5**). The PCA analysis for all recorded neurons in both pathways showed that the mPFC→BLA neurons exhibit greater separation between social (M1) and nonsocial (O) stimuli than the mPFC→NAc neurons (**Fig. 5A**). This indicates that mPFC→BLA neurons display a more distinct coding of social and nonsocial interactions. We quantified the distance between the neuronal activity clusters for social and nonsocial stimuli, showing a significantly greater separation in the mPFC→BLA pathway compared to the mPFC→NAc pathway, supporting the notion that mPFC→BLA neurons are more attuned to distinguishing between these stimuli (**Fig. 5B**). We then focused specifically on center-ON neurons identified during the OFT. The PCA plots showed that the center-ON neurons in the mPFC→NAc pathway exhibited more distinct separation between social and nonsocial stimuli compared to the mPFC→BLA neurons which showed much less differentiation (**Fig. 5C**). We further quantified the distances between social and nonsocial stimuli for center-ON neurons, with mPFC→NAc neurons showing significantly greater separation compared to mPFC→BLA neurons (**Fig. 5D**). Overall, the results demonstrate that while mPFC→BLA neurons exhibit broader population-level encoding of social versus nonsocial stimuli, the mPFC→NAc pathway is more specialized in differentiating these stimuli within behaviorally relevant neuronal subsets, such as center-ON neurons. This distinction highlights complementary roles for these pathways: mPFC→BLA neurons contribute to general contextual or environmental differentiation, whereas mPFC→NAc neurons are more specifically involved in encoding social preference.

**Figure 5.**
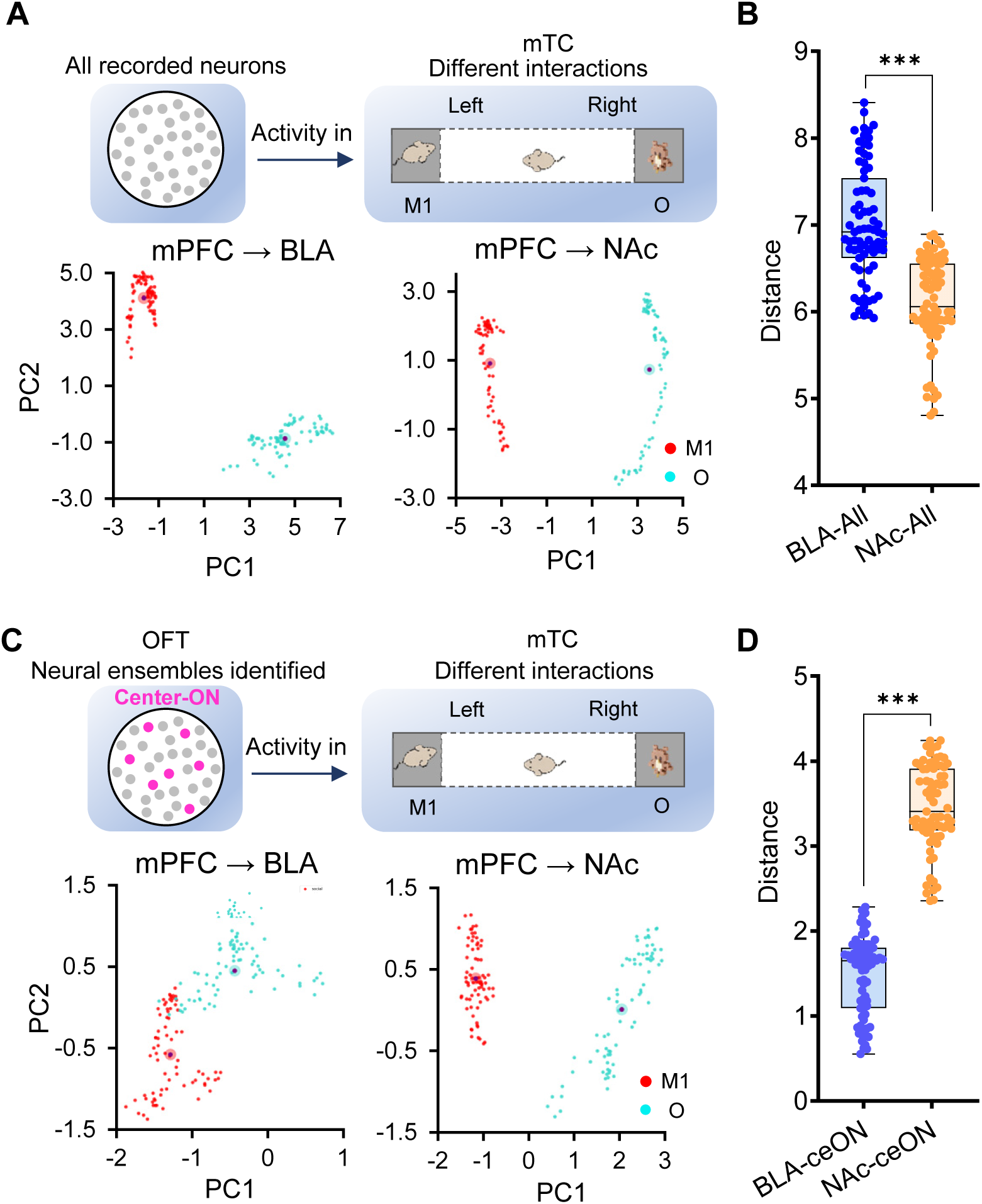
Comparative analysis of all neurons and center-ON subsets reflects divergent encoding patterns in the mPFC→BLA and mPFC→NAc Pathways. (A) PCA plots showing the activity of all recorded neurons in the mPFC→BLA and mPFC→NAc pathways during interactions with a social stimulus (mouse, M1) and a nonsocial stimulus (object, O) in a modified Three Chamber Test (mTC). The plots reveal distinct clustering of neural activity patterns for each stimulus within both pathways. (B) Distances between all neuronal activity clusters for nonsocial (O) interactions from the center of that for M1 interaction, compared across mPFC→BLA and mPFC→NAc pathways. Significant difference is demonstrated in the mPFC→BLA pathway. (C) PCA plots for center-ON neurons identified during the Open Field Test (OFT). These plots illustrate the activity of these neurons in the mPFC→BLA and mPFC→NAc pathways during the same social and nonsocial stimuli. Clusters show how Center-ON neurons specifically respond to each type of stimulus. (D) Distances of all center-ON neuronal activity clusters from the center of that for M1 interaction, compared across mPFC→BLA and mPFC→NAc pathways. Significant difference is demonstrated in the mPFC→NAc pathway. Data are represented as mean ± SEM. ***P < 0.001, Mann-Whitney U test.

To evaluate the consistency of our neuron co-registration across behavioral tests, we quantified the proportion of neurons reliably matched between sessions using the open-field test (OFT) as the reference. For the mPFC→BLA pathway, the mean yield of matched neurons was approximately 30% between OFT and EPM (range: 5-92%, n = 19 mice) and 16% between OFT and Social tests (range: 5-32%, n = 14 mice). For the mPFC→NAc pathway, the mean yields were 23% (range: 11-71%, n = 20 mice) and 21% (range: 13-37%, n = 17 mice) for OFT-to-EPM and OFT-to-Social alignments, respectively. Despite variability across animals, these values indicate that a substantial subset of neurons can be reliably tracked across days and behavioral contexts, supporting the stability of our longitudinal imaging alignment method.

### The mPFC→NAc and the mPFC→BLA neurons were differentially modulated by chronically induced affective states

Having established pathway-specific neural representations of behaviorally defined states (Figs. 1-5), we next asked whether these states correspond to biologically meaningful internal conditions. A key challenge in interpreting neural activity during behavior is determining whether such states reflect genuine internal processes rather than purely descriptive behavioral categories. To address this, we employed a repeated social competition paradigm (tube test) to establish stable social hierarchy and examine whether hierarchy-dependent differences are accompanied by measurable physiological and behavioral changes. Social competition outcomes, such as winning or losing, are ecologically valid means of eliciting opposing emotional experiences in rodents. The social dominance tube test has been widely used to assess social hierarchy and aggression (*22, 23*), a critical component of social stress and emotion regulation in rodents. It may therefore serve as an effective model for natural emotion induction. Studies have shown that social defeat or subordination can lead to measurable changes in neural activity, stress hormone levels, and behavior (*24–27*). Therefore, we believe that while the tube test may not induce long-lasting emotional changes on its own, it can serve as an effective and ethologically relevant model for natural emotion induction, particularly in the context of transient emotional responses associated with social competition and status.

In this assay, two mice are placed at opposite ends of a narrow transparent tube and allowed to interact as they attempt to pass through to the other side. (*22, 23, 28, 29*). Repeated daily over a week, this procedure often results in mice adopting consistent winner or loser roles, potentially leading to sustained positively or negatively valenced affective states (**Fig. 6A**). To control for potential time-dependent or nonspecific effects, a group of singly housed mice that did not undergo tube testing but were handled under identical conditions and on the same experimental timeline served as controls. Corticosterone levels measured from brain lysates revealed that loser mice exhibited significantly higher concentrations than both control and winner groups, indicating that the observed physiological changes were specific to social subordination rather than general effects of time or repeated exposure (**Fig. 6B**).

**Figure 6.**
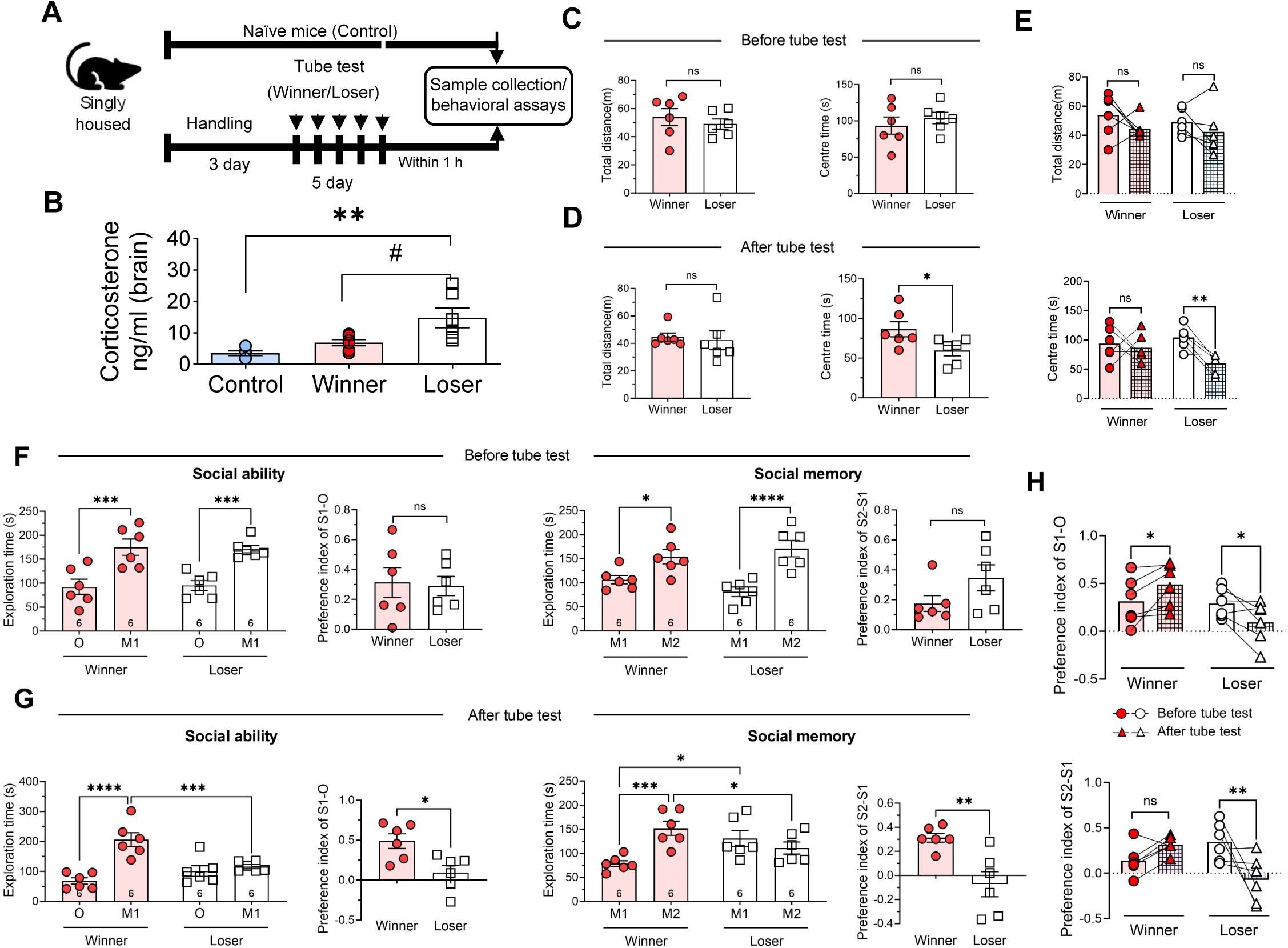
The modifications in social ranking of mice alter their anxiety and social states. (A) Schematic of the experimental timeline. Singly housed mice were handled for 3 days prior to undergoing a 5-day tube test to establish dominant (Winner) and subordinate (Loser) social status. Naïve singly housed mice served as controls. Behavioral assays were conducted 24 hours after, and brain tissue collection within 1 hour of, the final tube test session. (B) Corticosterone concentrations measured from brain lysates of Control, Winner, and Loser mice. Loser mice showed significantly elevated corticosterone levels compared to both Control and Winner groups (** P < 0.01, # P < 0.05). Data are presented as mean ± SEM. (C-D) The mice’s total distance traveled in the open field and time spent in the center zones before and after the tube test. *P < 0.05; ns, non-significant. n = 6 mice for each group, winner vs loser, unpaired Student’s t-test. (E) The differences of total distance (top) and center time (below) between winner and loser groups before and after the tube tests. **P < 0.01; ns, non-significant. Before vs. after tube test, Two-way ANOVA Multiple comparisons, sidak’s post-hoc test. (F) The mice’s exploration time near to M1 and O/M2 during the social ability test (left) and social memory test (right) conducted before the tube test. Social ability index calculated as (time near M1 chamber - time near object chamber) / (time near M1 chamber + time near object chamber). Social memory index calculated as (time near M2 chamber - time near M1 chamber) / (time near M2 chamber + time near M1 chamber). *P < 0.05, ***P < 0.001, ****P < 0.0001, ns, non-significant. Two-way ANOVA Multiple comparisons, sidak’s post-hoc test. Unpaired Student’s t-test for preference index. (G) The mice’s exploration time spent near the M1 and O/M2 during social ability test and social memory test after the tube test. *P < 0.05, **P < 0.01, ***P < 0.001, ****P < 0.0001. Two-way ANOVA Multiple comparisons, sidak’s post-hoc test. Unpaired Student’s t-test for preference index. (H) The comparisons of the social ability index (top) and social memory index (below) of winner and loser group before and after the tube tests. *P < 0.05, **P < 0.01, ns, non-significant. Two-way ANOVA Multiple comparisons, sidak’s post-hoc test). n = 6 mice for each group.

To verify if the tube test genuinely altered emotional states, littermate mice underwent an open field test both before and after the tube test (**Fig. 6C-E**). Initially, there was no difference in the travel distance or time spent in the center of the open field arena between winner and loser mice (**Fig. 6C**). Post-tube test, travel distances remained unchanged, suggesting that mobility was not affected by induced emotions (**Fig. 6D and E**). However, loser mice spent significantly less time in the center, indicating heightened anxiety and successful negative emotion induction. In contrast, winners’ time in the center remained unchanged, implying their anxiety levels remained normal (**Fig. 6D and E**). Furthermore, to determine if the tube test could evoke positive emotions in winner mice, their sociability was assessed before and after the test because positive emotion (such as the pleasure from winning) would enhance sociability. Initially, both the winner and loser mice showed a standard preference for interacting with a live mouse over an inanimate object, and a new mouse over a familiar one (**Fig. 6F**). After the tube test, winners displayed a higher sociability index towards social mice and a tendency for greater interest in new mice. In contrast, losers almost entirely lost their social preference, underscoring the induction of negative emotions (**Fig. 6G and H**). Together, these findings demonstrate that repeated tube test competition induces distinct, hierarchy-dependent behavioral and physiological changes. Importantly, these results provide independent validation that the behavioral states examined in earlier figures are associated with measurable internal changes, supporting their interpretation as meaningful proxies for affective state in the analysis of mPFC circuit dynamics. We note that the tube test is not a classical chronic stress paradigm and therefore interpret these effects conservatively as hierarchy-associated state changes rather than sustained emotional states.

## DISCUSSION

This study investigates the distinct roles of mPFC projections to the BLA and NAc in processing affective and social stimuli. Using a combination of behavioral paradigms and in vivo neural recordings, we found that although both pathways contribute to the encoding of emotionally salient and socially relevant behaviors, they exhibit divergent activity dynamics and functional specializations. We interpret the observed behaviors across the OFT, EPM, and tube test as validated proxies for internal emotional states, based on extensive prior literature. For example, avoidance of the center area in the open field test and the open arms in the elevated plus maze are widely accepted as behavioral indices of anxiety-like states in rodents (*14, 16*). Similarly, outcomes of the tube test have been shown to predict social hierarchy and are associated with long-lasting emotional valence shifts, including increased stress vulnerability in subordinate (loser) animals (*22, 23*). Prior research has shown that transitions between exploratory and defensive behaviors are associated with rapid shifts in neural ensemble activity in the BLA, consistent with changes in internal affective or motivational states (*30, 31*). These studies support the use of well-defined behavioral states as reliable proxies for emotional dynamics. While direct measurement of physiological stress markers such as corticosterone would further strengthen this link, such measurements are not feasible during in vivo calcium imaging, particularly during rapid state transitions in freely moving mice. Therefore, our interpretation of circuit activity in relation to “emotional states” relies on a validated behavioral framework supported by ensemble-level neural correlates.

The mPFC→BLA pathway demonstrated a broader capacity to differentiate between a range of affective states, particularly those associated with anxiety-like behavior, suggesting a generalized role in emotional processing of aversive or negatively valenced stimuli. In contrast, the mPFC→NAc pathway, especially its center-ON neurons, was more selectively tuned to exploratory behaviors and social interactions, indicating a role in encoding reward-related information and approach-oriented behavioral states (*6*). This differentiation is most evident in the context of anxiety-like behavior, where the mPFC→BLA neurons show enhanced responsiveness, highlighting their pivotal role in regulating anxiety-related states. This observation concurs with prior research highlighting the pivotal role of the Amygdala in the processing of fear and anxiety (*32, 33*).

The contrasting patterns of neuronal activity observed between “center-ON” neurons and the entire neuronal population further underscore the functional diversity within the mPFC→BLA and mPFC→NAc pathways (**Fig. 1 and 2**). The broader population of neurons does not exhibit the same selective sensitivity to affective state changes as the “center-ON” neurons. Instead, their activity appears more generalized, lacking the specificity observed in the center-ON subset. This suggests that while the overall activity within these pathways may contribute to the modulation of emotional responses, it is the center-ON neurons that provide a more precise and dynamic representation of affective shifts, particularly those related to anxiety-like behavior. Acting as a specialized subpopulation, center-ON neurons appear finely attuned to salient triggers such as entry into anxiogenic environments, offering a more detailed encoding of emotionally relevant contexts. In contrast, the broader neuronal population may support general regulation of affective states, albeit in a less differentiated manner. Together, these findings highlight the complexity of emotional processing within mPFC circuits, revealing the coexistence of specialized and generalized encoding mechanisms that collectively shape nuanced behavioral responses to emotionally salient stimuli.

Based on our findings in the social behavior paradigm, a more nuanced picture emerges of how different neuronal subsets within the mPFC pathways respond to social and nonsocial stimuli. At the population level, the mPFC→BLA pathway shows significantly greater differentiation between social and nonsocial stimuli compared to the mPFC→NAc pathway (**Fig. 5A, B**). This suggests that mPFC→BLA neurons are broadly tuned to encode differences across diverse stimulus categories, supporting generalized responses to varied environmental contexts. Such a role may be particularly relevant because anxiety-like states can be triggered by many types of cues, whether social or nonsocial, requiring a more holistic mode of neural processing. In contrast, within the center-ON neuronal subset, the mPFC→NAc pathway exhibits significantly greater differentiation between social and nonsocial stimuli compared to the mPFC→BLA pathway (**Fig. 5C, D**). This finding suggests that this subset of mPFC→NAc neurons is more specialized in encoding motivationally salient social cues, potentially playing a critical role in shaping specific aspects of social behavior. Importantly, while social preference is often associated with positive valence, the sharper tuning observed in mPFC→NAc neurons should be interpreted as enhanced encoding of social preference itself, with implications for, but not a direct measure of, affective valence. Together, these results highlight a layered encoding scheme in which the mPFC→BLA pathway provides a broad, generalized framework for environmental differentiation, whereas the center-ON neurons in the mPFC→NAc pathway selectively refine distinctions between social and nonsocial stimuli. This dual strategy may enable both flexible adaptation to general environmental contexts and precise behavioral responses in socially nuanced situations.

The current study focused on decoding the hidden variabilities in mPFC efferent pathways across emotional states of adult female mice, given that females are nearly twice as likely as males to be diagnosed with anxiety and depressive disorders (*34, 35*). Previous work (*36*) demonstrated that female rodents exhibit a higher density of dendritic spines on infralimbic mPFC neurons projecting to the BLA following fear conditioning, and show more rapid extinction of conditioned fear responses, suggesting structural and functional adaptations that differ from males. Stress responses differ between genders, with men showing increased activity in the prefrontal cortex and women in the limbic system (*37*). In female rodents, the neuronal firing activity of BLA shifts across the estrous cycle, which contributes to the sex-differences of BLA activity changes effected by repeated restraint stress treatment (*38*). Meanwhile, when receiving chronic variable stress, the functional and morphological changes of NAc-projecting mPFC neurons were more severe in female than in male mice (*39*). Finally, sex-specific transcriptional signatures occur in the PFC neurons of human patients with major depressive disorder (MDD), with females showing downregulation of *DUSP6* gene associated with increasing pERK in pyramidal neurons, which impacts on excitatory synaptic transmission in PFC of depressed female or stressed mice. On the other hand, upregulation of *EMX1* gene in PFC of male with MDD may increase their neuronal excitability and stress susceptibility (*40*). Taken together, these studies highlight pronounced sex differences in the structure, plasticity, and functional output of mPFC-BLA and mPFC-NAc circuits, which likely underlie the female-predominant prevalence of anxiety and depressive disorders. One limitation of the present study is that the experiments were performed primarily in female mice. Sex differences in prefrontal-limbic circuits have been reported in several behavioral paradigms, and future studies will be needed to determine whether similar projection-specific coding principles extend to male animals. Investigating how mPFC efferent pathways dynamically code emotional states in male animals will help elucidate the circuit-level mechanisms of sexually dimorphic behaviors.

It is also important to recognize that the diversity of activity observed here may, in part, reflect underlying anatomical and cellular heterogeneity. The mPFC→NAc pathway comprises neurons from multiple cell classes, and mPFC→BLA projections traverse different cortical layers. Likewise, both the BLA (magnocellular vs. parvocellular subdivisions) and the NAc (core vs. shell) are themselves heterogeneous structures with potentially distinct functional roles (*41, 42*). Future studies using more refined approaches will be needed to resolve how these subdivisions contribute to the pathway- and state-specific activity patterns identified in this work.

A central challenge in interpreting neural activity during behavior is establishing whether behaviorally defined states reflect meaningful internal conditions. In this study, we used commonly employed behavioral assays, including the open field test, elevated plus maze, and social interaction paradigms, to define distinct behavioral states associated with affective valence. Importantly, the tube test experiments provide independent validation that these states are associated with measurable physiological and behavioral changes, including altered corticosterone levels and shifts in exploratory and social behaviors. While the tube test does not constitute a classical chronic stress paradigm, these findings support the interpretation that the behavioral states used in our analyses capture biologically relevant internal conditions. This strengthens the link between mPFC circuit dynamics and affect-related behavioral states.

In summary, our study demonstrates that mPFC projection pathways make distinct contributions to behaviorally defined affective states in response to environmental cues. Neurons projecting to the BLA preferentially encode negatively valenced states and respond strongly to anxiogenic contexts, whereas those projecting to the NAc are more engaged during positively valenced states, including exploration and social preference. These findings advance our understanding of how prefrontal circuits organize behavior in response to environmental context and provide a framework for investigating how disruptions in these pathways may contribute to neuropsychiatric disorders. Importantly, these observations are correlational in nature. Establishing causal roles will require pathway- or ensemble-specific manipulations during behavior, an essential next step for directly linking neural dynamics to affective regulation.

## METHODS AND MATERIALS

Animal care and usage followed NIH Guidelines and received approval from the Institutional Animal Care and Use Committee of George Washington University and the University of Tennessee Health Science Center Laboratory Animal Care Unit.

### Experimental Animals

CaMKII-Cre mice with a pure C57BL/6 background were acquired from Jackson Lab (JAX#005359). For excitatory neuron imaging experiments, female CaMKII-Cre mice were bred by mating male CaMKII -Cre mice with female mice from the 129S1/SvlmJ strain (JAX#002448). These experimental CaMKII -Cre mice were used in accordance with the same procedural guidelines. Surgeries were performed on mice at approximately 4 months of age, adhering to the established experimental protocol. Animals were housed in groups of 4 to 5 per cage in a controlled environment with a temperature of 23 ± 1°C, humidity set at 50 ± 10%, and a 12-hour light-dark cycle. Standard mouse chow and water were provided ad libitum.

### Virus injection and gradient-index (GRIN) lens implantation

In the context of imaging excitatory neurons, we employed the AAV1-EF1a-flex-GCaMP6m virus acquired from Baylor College of Medicine. Following established protocols (*21*), CaMKII - Cre mice were anesthetized and securely positioned within a Neurostar stereotaxic frame from Tübingen, Germany. Employing a high-speed rotary stereotaxic drill (Model 1474, AgnTho’s AB, Lidingö, Sweden), we conducted a unilateral injection of the retrograde virus into either the left NAc region (anterior-posterior AP: +1.34 mm, medial-lateral ML: -1.1 mm, dorsal-ventral DV: 4.4mm) or the Amygdala region (AP: -1.6 mm, M/L: -2.9 mm, DV: 4.5mm) using the Nanojector II system from Drummond Scientific. The injection comprised 300 nL of virus diluted with 300 nL of phosphate buffer solution (PBS), administered at a controlled rate of 30 nL/min. Post-injection, the needle was retained in place for an additional 5 minutes to optimize virus diffusion.

Subsequent to viral injection, a precise 1.1 mm-diameter craniotomy was conducted at AP: +1.95 mm, M/L: -0.5 mm coordinates. These values were determined relative to bregma: +1.95 mm AP, -0.35 mm ML, -2.3 to -2.5 DV, employing a high-resolution atlas. Following this, a 1-mm diameter GRIN lens (Inscopix, Palo Alto, CA) was progressively lowered into the left PL region (AP: +1.95 mm; ML: ± 0.35 mm; DV: -2.1∼-2.3 mm) at a rate of 50 μm/min, situated 0.2 mm above the virus injection site, and then cemented in place using Metabond S380 (Parkell). Subsequently, the mice were allowed to recover on a heating pad and were closely monitored over the ensuing 7 days, during which they received analgesic treatment.

### Baseplate attachment

Around three to four weeks post-surgery, we assessed virus expression in the anesthetized mice using a miniaturized microscope from Inscopix, based in Palo Alto, CA. Once GCaMP+ neurons were clearly visible, we proceeded to attach the microscope with a baseplate onto the mouse’s skull window. This setup was then gradually lowered to determine the optimal focus plane. Subsequently, the baseplate was securely affixed to the skull using dental cement and covered with a protective cap, while the microscope remained unattached. Ahead of the behavioral tests, the mice were familiarized with the test room environment, during which a dummy microscope was mounted and handled for approximately 5 to 7 days, with sessions lasting 30 to 40 minutes each day.

### Selection of animals

Mice were chosen based on specific criteria: for the observation of Ca^2+^ signals in the mPFC, mice were excluded post-hoc if (1) the GRIN lens was positioned outside of the prelimbic cortex, (2) GCamp6 was not expressed within prelimbic areas, (3) significant virus expression was observed outside of the prelimbic region, and de(4) the imaging plane was obstructed by blood or debris. In the identification of mPFC neural ensembles, mice exhibiting no center entry behavior were excluded, as the classification was grounded in the neural responses to transitions in location.

### Determination of sample size

The sample sizes for behavioral experiments were established following the prevailing standards in behavioral neuroscience research for mice. This approach considers the minimum number of mice needed to detect statistical significance, with an α level of 0.05, and a statistical power of 80% or higher. Our post-hoc analysis demonstrates a statistical power of 85%.

In terms of neuron count, we recorded 11-83 neurons expressing GCaMP6m for the mPFC→BLA group (with an average of 38) and 18-54 for the mPFC→NAc group (with an average of 39). This range was influenced by various factors, including the injected volume of viral GCaMP6m, expression levels, and the efficiency of Cre-mediated recombination. Following the data processing phase, which involved the identification of neurons through principal component and independent component analyses (PCA-ICA), approximately 1-9% of the identified components were identified as artifacts and subsequently excluded. The remaining components were classified as neurons and employed for subsequent analyses.

#### Behavioral tests

The behavioral tests spanned three consecutive days, maintaining consistent schedules throughout. Behavioral assays were separated by at least 24 hours to minimize potential effects of fatigue or prior experience. To ensure cleanliness, the chamber was meticulously cleaned with 70% ethanol between trials. The Topscan behavior analysis system from Clever Sys, VA, was employed to monitor the animals’ behaviors. This system also sent a TTL signal at the start of each test to trigger the microscope recording of neuronal activity.

### Modified Three-Chamber (mTC) Test

The mTC test adhered to a previously established protocol (*43*) with several enhancements. The conventional three-chamber apparatus was transformed into a single open box (45 x 10 x 20 cm) featuring two small removable lateral chambers (10 x 10 x 40 cm). These chambers were separated by thin, spaced metal wires, permitting mice to interact with stimuli. Before the test, the subject mouse underwent a 10-minute habituation period within the open box devoid of stimuli. During testing, an age- and weight-matched, unfamiliar same-sex conspecific (referred to as the first social stimulus, M1) and an inanimate object (non-social stimulus, O) were randomly placed into the two lateral chambers. Subsequently, the subject mouse was positioned in the center of the open box and allowed to explore freely during a 10-minute testing session. Several behavioral parameters were assessed, including time spent in social interaction, object interaction, the social zone (SZ), the object zone (OZ), the transition zone, and grooming behavior.

### Open field test (OFT)

The test was modified from a previous report (*14*). It was conducted in a square box (dimensions: 50 × 50 × 50 cm). The mouse was gently placed in the central field and allowed to explore freely during a 10-minute testing session. Locomotor activity was recorded by a camera. The center is defined as the central 25 cm x 25 cm square area, while the corner is a sector area with a 12.5 cm radius in each corner. Total distance traveled and time spent in each area, sniffing, and grooming behaviors were analyzed using the TopScan automated behavioral analysis system (CleverSys Inc., Reston, VA). “Sniffing” behavior was defined as episodes when the mouse’s nose was within approximately 1 cm of the wall and moving in small scanning motions without locomotion away from the wall, lasting for at least 1 second. To ensure clear separation from “corner” behavior, sniffing events occurring within the defined corner zones were excluded from the corner category and analyzed separately. As a result, corner time does not include sniffing episodes that occurred in corners.

### Elevated plus maze (EPM) test

It was conducted as previously described (*16*). The EPM apparatus, 40 cm high from the floor, consists of two open arms (35 × 10 cm) and two closed arms (35 × 10 cm), those two parts stretch perpendicular to each other and connect to a center platform (5 cm). Mice were placed in the center zone facing an open arm and allowed to explore the maze freely for 10 min. The time spent in open arms, closed arms, and behaviors of head dipping, sniffing, and grooming were analyzed.

### Dominance tube test

The tube test protocol was modified from Wang et al (*23*). In brief, a clear acrylic tube with a 30 cm length and 2.8 cm inside diameter in which one mouse can pass or backward through the tube fluently, but cannot make a U-turn or climb through another mouse. The mice underwent a three-day training in which all mice went through the tube ten times per day, five times from each side, without another competitor in the tube. The training procedure allows the mice to adapt to the test procedure and environment. The mice, in a very rare case, cannot be trained and were excluded from further testing. In the testing procedure, tests were conducted in a pair-wise style, and the number of times won by individual animals was recorded to determine the hierarchical ranking. The mouse that forced its competitor out of the tube was declared the “winner”, or dominant in this situation. The mouse that was retreated is then designated the “loser” or subordinate. In most cases, the competition was completed within 2 minutes, or the tests were repeated. The rank is considered stable without a ranking change in all mice for at least five consecutive days. Groups were categorized as “non-stable ranking” if they did not form a stable ranking within 2 weeks. We single-housed the mice for 5 days and subjected them to a round-robin tube test tournament (three trials per day for 7 trials) to determine social ranking. Twenty-four hours following the last tube test, the mice were subjected to other behavioral tests or in vitro studies, as described in the results.

### Behavior data analysis

Behavioral data was meticulously tracked through aerial videography from an overhead perspective utilizing the Topscan behavioral data acquisition software (CleverSys, Reston, VA). This software allows for the precise tracking and definition of the 2D locations of mice in various areas, including the OF center, corner area, EPM open-P, open-NP, closed-P, closed-NP arms, and mTC social zone, object zone, and middle zone. Additionally, specific behaviors such as sniffing, grooming, head-dipping, and social/object interaction were recognized and quantified by the software.

#### Ca^2+^ imaging with Miniature microscope

Imaging of freely moving mice was conducted using a head-mounted miniaturized microscope (nVista HD 2.0.4, Inscopix, Palo Alto, CA), as depicted in **Fig. 1A**. This microscope was synchronized with the Topscan system through a TTL pulse, enabling simultaneous acquisition of Ca^2+^ signals and behavioral video. Prior to imaging, the microscope was securely affixed to the mouse’s head. The imaging data were obtained at a frame rate of 15 Hz with a resolution of 1024 x 1024 pixels. LED power settings ranged from 0.3 to 1 mW, while gain settings were adjusted to 1 to 2 based on fluorescence intensity. Importantly, each individual mouse utilized the same imaging parameters consistently across all three experimental sessions.

#### Histology

Recording sites were meticulously validated through histological examination of lesions created during the lens implantation procedure. Mice were anesthetized via intraperitoneal injection, employing a combination of ketamine (400 mg/kg) and xylazine (20 mg/kg). Following this, transcardial perfusion was conducted using PBS, followed by 4% paraformaldehyde (PFA). The brains of the mice, complete with their skulls and baseplates, were post-fixed with 4% PFA for 3 days. Subsequently, the brains were extracted and sectioned into slices measuring 50-100 µm using a vibrating slicer (Vibratome Series 1000, St. Louis, MO). These sections were then mounted onto slides. To label cell nuclei, slides were subjected to incubation and storage in a 1:1000 Hoechst solution in 1x PBS (Invitrogen, Carlsbad, CA). Brain slides were meticulously imaged to precisely determine the placement of the GRIN lens and the extent of viral expression. This imaging process was conducted employing a Confocal Microscope (Zeiss LSM 710, Oberkochen, Germany).

#### Ca^2+^ image processing

Ca^2+^ images were processed offline using Inscopix Data Processing software (version 1.3.1). In brief, frames collected within a single day were concatenated into a stack and subsequently subjected to preprocessing, spatial filtering, and motion correction. To normalize the Ca^2+^ signal, the average projection of the filtered video was established as the background fluorescence (F_0_). Instantaneous normalized Ca^2+^ fluorescent signals (ΔF/F) was calculated according to the formula, (ΔF/F)_i_=(F_i_-F_0_)/F_0_, where i represents each frame. Individual cells were then identified using the principal component and independent component (PCA-ICA) analyses with no spatial or temporal down-sampling. Regions of interest (ROI) were selected based on signal and image criteria, with any components that did not correspond to single neurons being discarded.

Time-stamped traces of neurons were exported into data files formatted for custom-written Python scripts used for subsequent analysis. Ca^2+^ transients (events) were identified for each cell using a peak-finding algorithm, and frequency (transient rate) and amplitude (ΔF/F) data were processed using custom-written scripts. When analyzing frequency or amplitude changes in response to mouse behavior or location, frames of image and behavioral data were aligned and marked with corresponding behavioral event labels. For neuronal activity normalization, we employed the min-max normalization method to scale values between 0 and 1. In this method, the maximum neuron activity value was transformed into 1, while all other values were scaled to decimals between 0 and 1 using the equation: 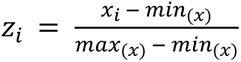, where x = (x1,x2,…,xn) represents an array of neuron frequency or amplitude values, and z_i_ denotes the normalized value.

### Identification of location-modulated neural ensembles

To identify location-modulated neural ensembles in each session, we assessed each neuron’s response preference to a specific location exploration. Initially, we computed the actual similarity (Sa) between the Ca^2+^ trace vectors (c_k_) and behavior events (b), using the formula: 2b ⋅ c_k_/(|b|^2^ + |c_k_|^2^) (*18*). Then, the behavior vector was subjected to random shuffling to compute a new similarity (S) with a neural trace for a given neuron. This shuffling process was repeated 5000 times, generating an S distribution histogram. Neurons were classified as ON neurons if their Sa values exceeded the 99.95th percentile of the S distribution.

Following neuron identification, we calculated the proportions of different location-modulated ON neurons for each mouse. Simultaneously, the transient rate and amplitude of these neuron ensembles were computed for each mouse in each paradigm. Animals in which no neurons met the predefined criteria for center-ON classification were excluded from analyses specifically requiring this classification, as inclusion would preclude meaningful comparison of center-ON population activity.

### Neuron alignment between tests

To identify and compare the same neurons across tests conducted on different days, thus enabling cross-test comparisons, we implemented a global alignment procedure. Initially, we reconstructed neuron distribution images based on their pixel locations and shapes.

Subsequently, we converted each neuron image into a vector format and assessed its similarity using cosine similarity, as indicated by the formula below. The two parameters, degree of rotation (A) and pixel shift (B), were iteratively adjusted to maximize the cosine similarity of image vectors between tests. Given that the images from all three tests were collected under identical settings, the scale for the neuron images remained constant and did not require adjustment. Through this multi-start method, we achieved the global optimal solution for the parameters. Ultimately, we determined the number of overlapping neuron images across tests, which was then used for subsequent comparative analyses.

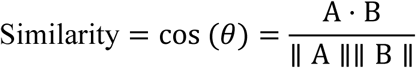

### Principal Component Analysis (PCA)

Population neural activity was analyzed using principal component analysis (PCA) to reduce dimensionality and visualize population dynamics across behavioral states. Calcium event rates were z-scored for each neuron across time before concatenation across mice to generate a population activity matrix (frames × neurons) for each pathway. PCA was applied to this pooled matrix using the covariance method implemented in MATLAB (MathWorks) and scikit-learn (Python), centering each neuron’s activity by its mean to project neurons from different animals into a common coordinate space.

The first three principal components (PCs) were used to compute neural trajectories and distance metrics, as they captured the majority of the variance in population activity (∼49% for all neurons and ∼59% for center-ON neurons in the mPFC→BLA pathway). To quantify neural representation differences across behaviors, we calculated the Euclidean distance from each behavioral cluster to the Corner cluster within the same PC space. Corner behavior was chosen as the reference because mice spent most of their time in the corners during the open-field test, representing a stable low-anxiety baseline state. Euclidean distances between behavioral clusters were computed using the centroid coordinates of each behavior (Center, Sniffing, Grooming, and Corner) in the two-dimensional PCA space defined by the first two principal components (PC1 and PC2), averaged across mice for each pathway.

### Cross-animal analysis

For cross-animal population analyses (e.g., **Fig. 1I**), neuronal calcium activity from all mice was concatenated along the neuron dimension to create a population matrix (frames × neurons). PCA was applied directly to this pooled dataset. Because PCA centers each neuron’s activity by its mean, neurons from different animals were projected into a shared coordinate frame without requiring further normalization. This approach enabled consistent visualization of behavioral state trajectories across animals in a common PC space.

#### Measurement of corticosterone levels in mPFC tissue

Corticosterone concentrations in the mPFC were quantified using a commercially available Cortisol ELISA kit (IBL America, IB79175) following the manufacturer’s instructions, with minor modifications for tissue samples. Briefly, mPFC tissues were rapidly dissected on ice and weighed using a micro-scale. 500μl of Ethyl Acetate was added onto the samples prior to homogenization on ice with a hand-held homogenizer: 10 seconds on and 10 seconds off 4 times. An additional 500μl of Ethyl Acetate was added onto the samples and then thoroughly vortexed (∼2min) and centrifuged at 500 g for 5 min at 4°C and left on dry ice for at least 6 hours. Supernatant was transferred to a new 2 ml Eppendorf tube and dried using a SpeedyVac rotary vacuum evaporator at 50°C. Samples were then stored at -80°C until used. Before performing the assay, samples were resuspended in 25 μl of 100% Ethanol. Resuspension of the samples in 0 calibrator or PBS resulted in the need for a large volume, reducing the concentration of corticosterone below the detection limit. The ELISA assay was performed following the manufacturer’s instructions, except that the assay buffer was added first, and 10μl of the samples were added on top of that. This is to avoid direct exposure of the antibodies to ethanol. Absorbance was then measured at 450 nm using an EnVision plate reader. The concentrations were calculated from a standard curve and normalized to total protein content determined by the BCA assay.

#### Statistics

All statistical analyses were performed using SPSS (version 24, IBM, Armonk, NY), Microsoft Excel (Redmond, WA), and custom Python scripts. Data were first tested for normality using the D’Agostino-Pearson test. For comparisons between two dependent conditions (e.g., data obtained from the same mouse), a two-tailed paired t-test was applied; for independent two-group comparisons, an unpaired two-tailed t-test was used. When comparing more than two groups, one-way ANOVA followed by Tukey’s post hoc multiple-comparison test was conducted. For datasets involving two experimental factors, two-way mixed-effects ANOVA with Bonferroni-corrected post hoc tests was performed. Statistical significance was defined as p < 0.05 (p < 0.01, highly significant), and non-significant results (e.g., p > 0.05 in paired t-tests) are reported where relevant. Normality was assessed using the Shapiro-Wilk test. When normality assumptions were violated or sample sizes were small (n < 15), non-parametric Mann-Whitney U tests were additionally performed. Effect sizes were reported as Cohen’s d for parametric tests and rank-biserial correlation for non-parametric tests. All data are presented as mean ± SEM, unless otherwise indicated.

To validate the significance of distance metrics in neural state space, permutation-based null controls were implemented by (i) shuffling behavioral labels within each mouse and session while preserving bout counts and trajectory lengths, and (ii) applying circular time-shifts per neuron within bouts to maintain each neuron’s autocorrelation. Observed cluster distances and pathway differences were compared against 10,000 permutation-derived null distributions, with significance defined as exceeding the 95th percentile (p_perm < 0.05).

## DATA AVAILABILITY

All data that support the findings in this study and the code used to analyze the data are available upon reasonable request.

## ACKNOWLEDGEMENTS

This work was supported by the NIH grant R01NS118197, and the George Washington University 2018-2023 Cross-Disciplinary Research Fund to H. L. and R.S. / C. Z. The National Institutes of Mental Health (R01MH135862), the Cystic Fibrosis Foundation (002544I221), and the University of Tennessee Health Science Center start-up fund to J.D.

## AUTHOR CONTRIBUTIONS

H.L. and J.D. conceived the project. H.L., J.D., C.L., G.P., P. X., Q.G., Z.J. designed the experiments. C.L. and P.X. performed the in vivo Ca^2+^ experiments. C.L., P.X., X.S., Y.L., R.S., and C.Z. performed imaging data analysis. Q.G. and Z.J. performed the behavior experiments. Z.J. and G.P. performed the mouse surgery. X.L. and Q.L. performed histology. C.L., H.L., and J.D. wrote the manuscript. All authors reviewed and edited the manuscript.

## ADDITIONAL INFORMATION

Supplementary information is available on eLife’s website.

---

**Supplemental Fig. 1.**
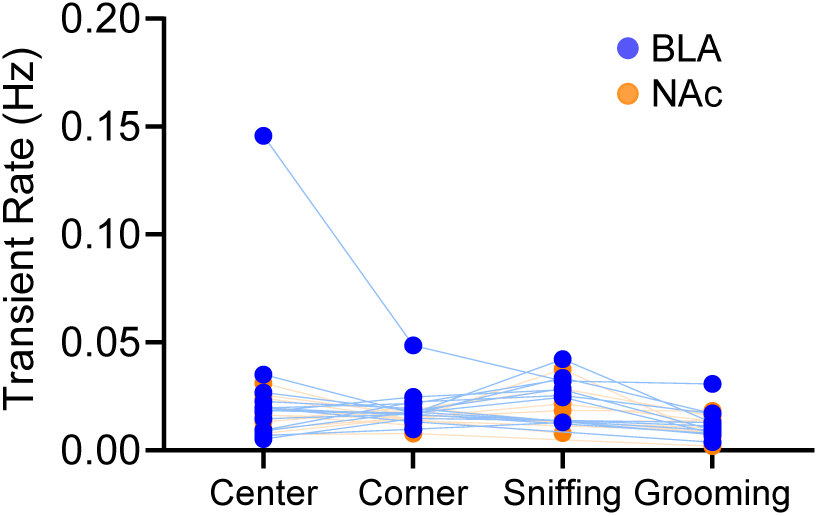
Individual data points for transient rate across behavioral states (related to Fig. 1H). Scatter plot showing transient rates of all mPFC→BLA (blue and mPFC→Nac (orange) neurons across four behavioral states (Center, Corner, Sniffing, and Grooming). Each blue dot represents data from an individual mouse, with connecting lines indicating within-animal comparisons across behaviors. The plot demonstrates that the observed group differences in Figure 1H are not driven by outliers, as individual data points show consistent trends across animals.

**Supplemental Fig. 2.**
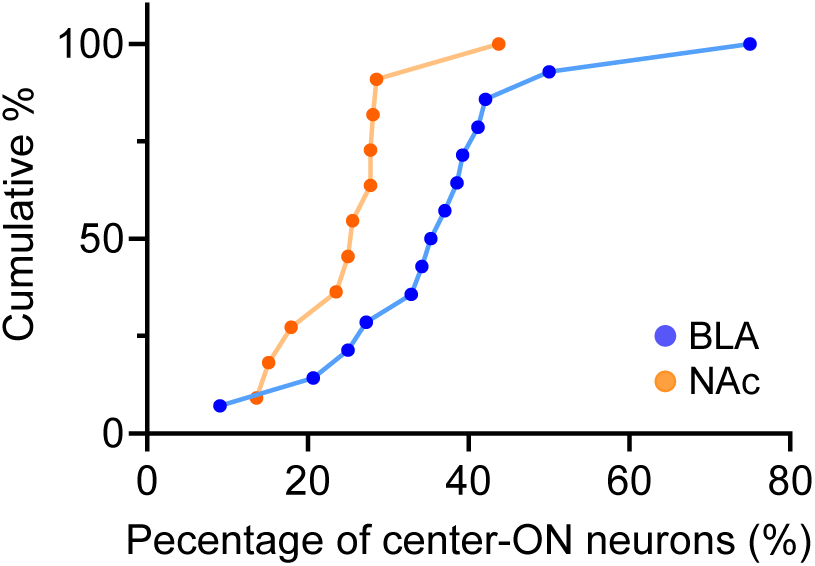
Cumulative distribution of center-ON neuron percentages across mice (related to Fig. 2C). Cumulative plots showing the percentage of center-ON neurons per mouse for the mPFC→BLA (blue) and mPFC→NAc (orange) pathways. The distribution of center-ON neuron percentages was significantly left-shifted in mPFC→NAc compared to mPFC→BLA neurons (two-sample Kolmogorov–Smirnov test, D = 0.61, p = 0.032), confirming a lower overall proportion of center-ON neurons in the mPFC→NAc pathway.

**Supplemental Fig. 3.**
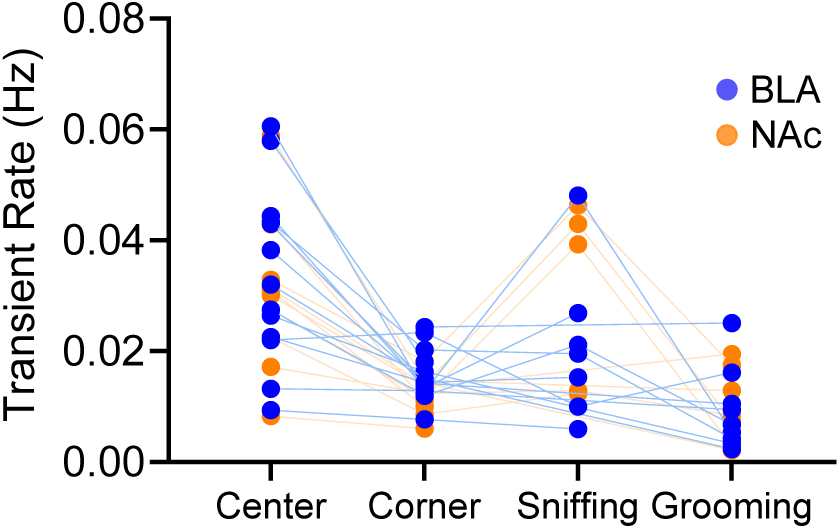
Individual data points for transient rate of center-ON neurons across behavioral states (related to Fig. 2E). Scatter plot showing transient rates of mPFC→BLA (blue) and mPFC→NAc (orange) center-ON neurons during Center, Corner, Sniffing, and Grooming behaviors. Each dot represents the average transient rate from an individual mouse, with connecting lines indicating within-animal comparisons across behavioral states. The plot confirms that the group differences shown in Figure 2E are consistent across animals and are not driven by outliers.

**Supplemental Fig. 4.**
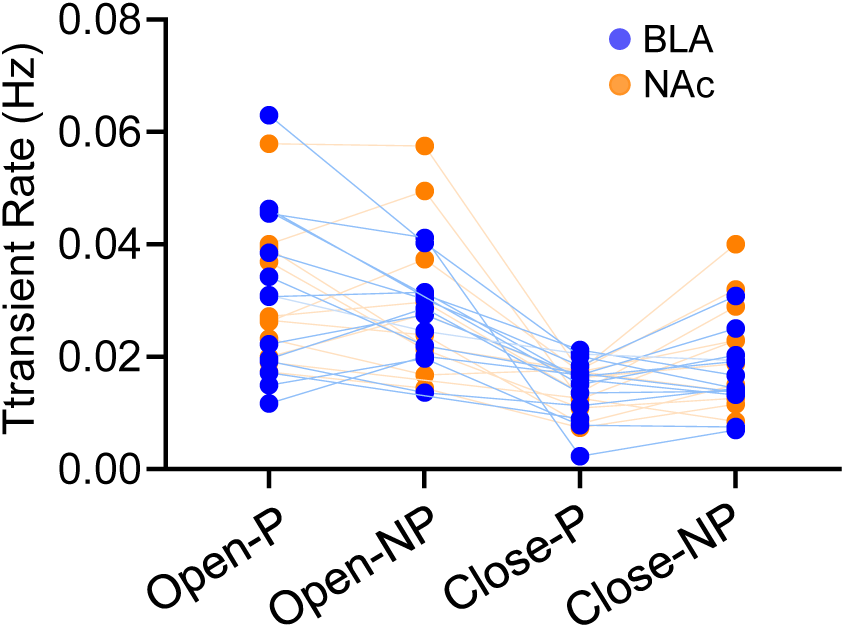
Individual data points for transient rates of all neurons in different arms of the elevated plus maze (related to Fig. 3C). Scatter plot showing transient rates of all recorded mPFC→BLA (blue) and mPFC→NAc (orange) neurons during Open Arm, Closed Arm, Center, and Grooming periods in the elevated plus maze (EPM). Each dot represents the mean transient rate from an individual mouse, with connecting lines indicating within-animal comparisons across behavioral conditions. The consistent distributions across animals confirm that the differences observed in Figure 3C are robust and not driven by individual outliers.

**Supplemental Fig. 5.**
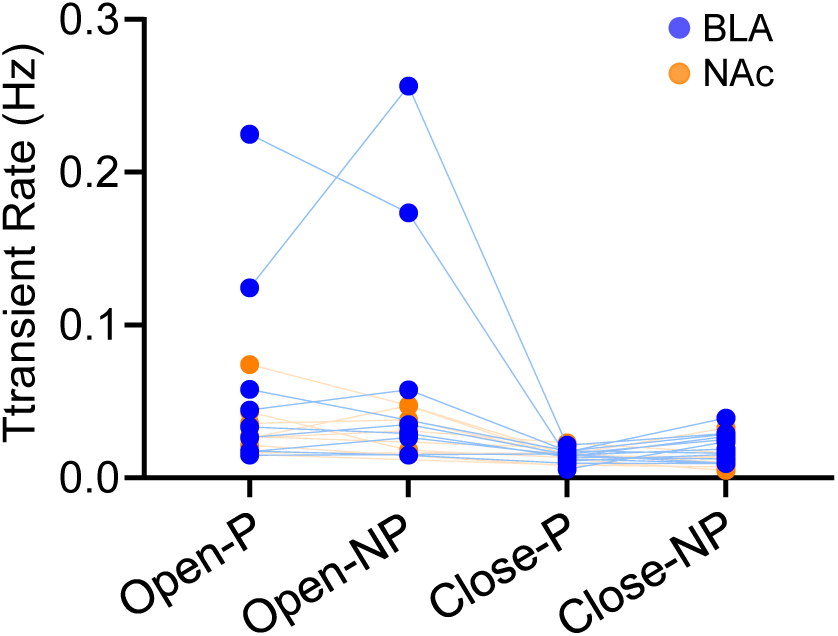
Individual data points for transient rate of in different arms of the elevated plus maze (EPM) (related to Fig. 3G). Scatter plot showing transient rates of mPFC→BLA (blue) and mPFC→NAc (orange) center-ON neurons of in different arms of EPM. Each dot represents the average transient rate from an individual mouse, with connecting lines indicating within-animal comparisons across behavioral states. The plot confirms that the group differences shown in Figure 3G are consistent across animals and are not driven by outliers.

